# Efficacy and dynamics of self-targeting CRISPR/Cas constructs for gene editing in the retina

**DOI:** 10.1101/243683

**Authors:** Fan Li, Sandy S.C. Hung, Jiang-Hui Wang, Vicki Chrysostomou, Vickie H.Y. Wong, James A. Bender, Leilei Tu, Alice Pébay, Anna E King, Anthony L. Cook, Raymond C.B. Wong, Bang V. Bui, Alex W. Hewitt, Guei-Sheung Liu

## Abstract

Safe delivery of CRISPR/Cas endonucleases remains one of the major barriers to the widespread application of *in vivo* genome editing including the anticipatory treatment of monogenic retinal diseases. We previously reported the utility of adeno-associated virus (AAV)-mediated CRISPR/Cas genome editing in the retina; however, with this type of viral delivery system, active endonucleases will remain in the retina for an extended period, making genotoxicity a significant consideration in clinical applications. To address this issue, we have designed a self-destructing “kamikaze” CRISPR/Cas system that disrupts the Cas enzyme itself following expression. Four guide RNAs (sgRNAs) were designed to target *Streptococcus pyogenes* Cas9 (SpCas9), after *in situ* validation, the selected sgRNAs were cloned into a dual AAV vector. One construct was used to deliver SpCas9 and the other delivered sgRNAs directed against SpCas9 and the target locus (yellow fluorescent protein, YFP), in the presence of mCherry. Both constructs were packaged into AAV2 vector and intravitreally administered in C57BL/6 and *Thy1-YFP* transgenic mice. After 8 weeks the expression of SpCas9, the efficacy of *YFP* gene disruption was quantified. A reduction of SpCas9 mRNA was found in retinas treated with AAV2-mediated-YFP/SpCas9 targeting CRISPR/Cas compared to those treated with YFP targeting CRISPR/Cas alone. We also show that AAV2-mediated delivery of YFP/SpCas9 targeting CRISPR/Cas significantly reduced the number of YFP fluorescent cells among mCherry-expressing cells (~85.5% reduction compared to LacZ/SpCas9 targeting CRISPR/Cas) in transfected retina of *Thy1-YFP* transgenic mice. In conclusion, our data suggest that a self-destructive “kamikaze” CRISPR/Cas system can be used as a robust tool for refined genome editing in the retina, without compromising on-target efficiency.

## INTRODUCTION

Inherited retinal diseases are disabling disorders of visual function and have affected millions of people worldwide. With the development of next-generation sequencing and better molecular diagnostic techniques, numerous genetic variants across many loci have been definitively associated with inherited retinal diseases.^1,2^ Despite this increase in our understanding of genetic aetiology and potential therapeutic targets, there remains no effective treatment for the majority of inherited retinal diseases.^3^ Although significant progress in gene therapy have been achieved over the last two decades, a sustained, safe and effective ocular gene therapy for hereditary retinal diseases is not readily available for all conditions.^4,5^

Genome editing techniques, in particular the recent advances in CRISPR/Cas technology, has renewed excitement in ocular gene-based therapy.^6^ The CRISPR/Cas system has evolved in archaea and bacteria as an adaptive defense against viral intrusion and has manipulated to allow efficient editing of mammalian nuclear genomes.^7^ CRISPR/Cas-based technology has proven a robust means for *in vitro* correction of genetic mutations in mammalian cells and is particularly attractive for treating inherited retinal diseases.^3^ A number of *in vivo* studies in various animal models have yielded promising results opening the prospect for preemptive therapy for well-characterised monogenic ocular diseases. Bakondi *et al*.^8^ and Latella *et al*.^9^ report successful ablation of the mutated rhodopsin gene prevented retinal degeneration in rodent models of severe autosomal dominant retinitis pigmentosa following electroporation of the CRISPR/Cas system into the retina. We were able to achieve high efficiency of genome editing in mouse retina using a dual AAV2-mediated CRISPR/Cas system.^10^ More recently, Yu *et al*.^11^ further demonstrated that CRISPR/Cas-mediated disruption of a neural retina-specific leucine zipper protein (*NRL*) significantly improved rod survival and preserved cone function in a murine model of retinal degeneration. Despite these promising applications in inherited retinal diseases, potentially deleterious off-target effects of CRISPR/Cas must be addressed, and it is well appreciated that prolonged over-expression of CRISPR/Cas endonucleases could result in elevated off-target cleavage,^12,13^ or potentially trigger cellular immune responses,^14^ thereby presenting important safety concern for clinical applications.

To address the potential for deleterious effects of CRISPR/Cas over-expression, we have designed a self-destructive “kamikaze” CRISPR/Cas system that disrupts the CRISPR/Cas gene after active protein expression (Figure 1). To determine the efficacy of *in vivo* genome editing by our “kamikaze” CRISPR/Cas construct, a SpCas9 targeting sgRNA module, together with a yellow fluorescent protein (*YFP*) targeting sgRNA, were packaged into a dual AAV2 vector system for intravitreal delivery in *Thy1-YFP* transgenic mice. Overall, our data demonstrates the feasibility of a self-destructive CRISPR/Cas system as a safe and robust tool for refined genome editing in the retina.

**Figure 1.**
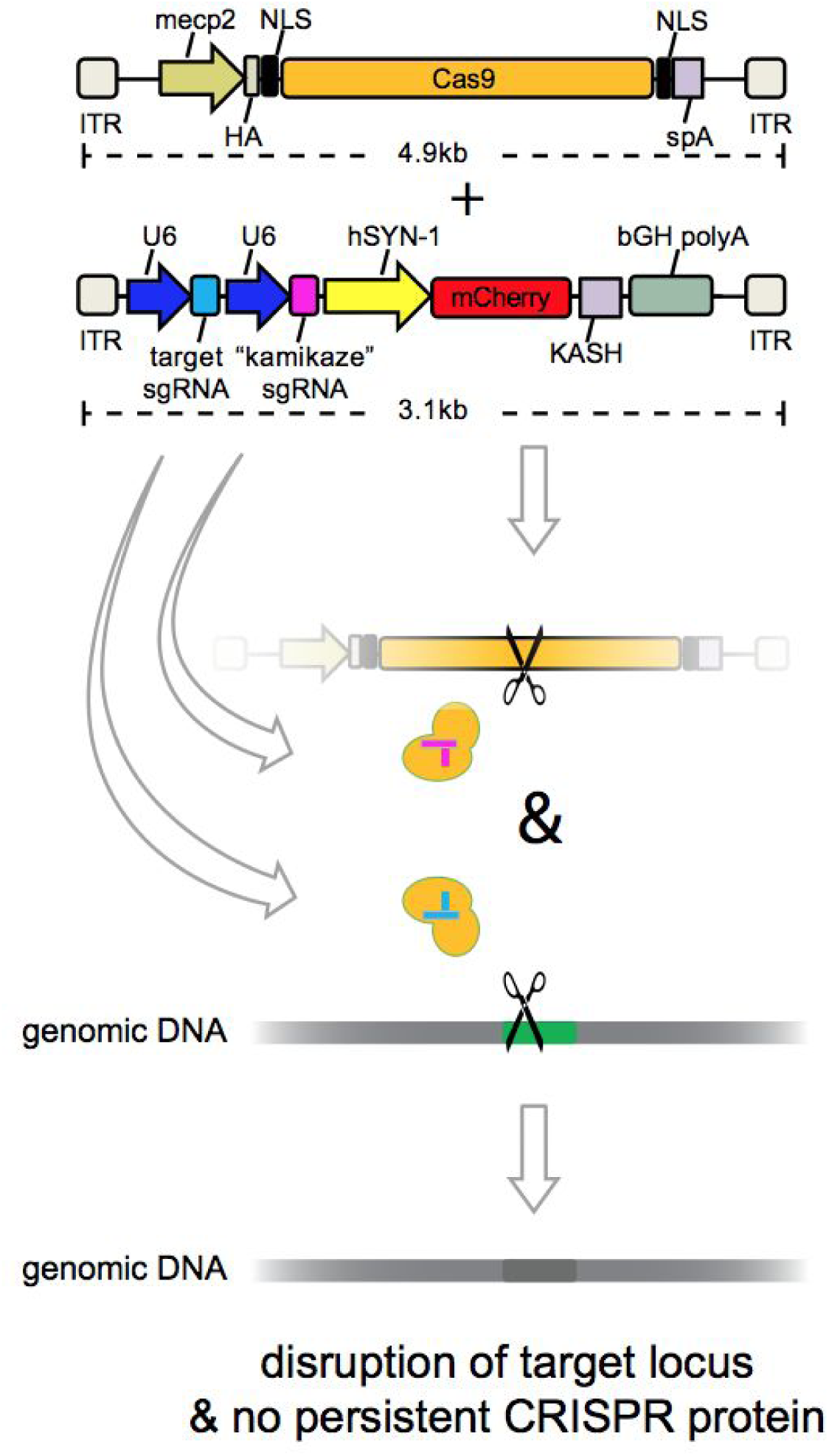
Schematics of Kamikaze CRISPR/Cas system. A dual AAV vector system was used. One viral vector was used to deliver SpCas9 and the other delivered sgRNAs against SpCas9 and the target locus (YFP), in the presence of mCherry.

## MATERIALS AND METHODS

### Animals and housing

*Thy1-YFP* transgenic mice [B6.Cg-Tg(Thy1-YFP)16Jrs/J] were obtained from the Jackson Laboratory (mouse stock number: 003709; Bar Harbor, ME, USA) and bred at the animal facility of the Menzies Institute for Medical Research (Hobart, TAS, Australia). C57BL/6 mice were purchased from the Animal Resources Centre (Perth, WA, Australia). Mice were housed under standard conditions (20°C, 12/12 hour light/dark cycle) with *ad libitum* access to food and water. All procedures were conducted according to the Association for Research in Vision and Ophthalmology Statement for the Use of Animals in Ophthalmic and Vision Research and the requirements of National Health and Medical Research Council of Australia (Australian Code of Practice for the Care and Use of Animals for Scientific Purposes). Ethics approval was obtained from Animal Ethics Committee of the University of Tasmania (A14827) and St. Vincent’s Hospital Melbourne (AEC 014/15).

### sgRNA design and vector construction

Four sgRNAs targeting the SpCas9 sequence were designed using a web-based CRISPR design tool (http://crispr.mit.edu). CRISPR/Cas *in situ* testing was carried out by incubating the individual synthetic SpCas9 sgRNA or LacZ sgRNA alone with recombinant SpCas9 protein (catalog no. M0386S; New England Biolabs, Ipswich, MA, USA) and the pX551 plasmid (SpCas9 construct; kindly provided by Feng Zhang, Addgene #60957). Samples were run on a 0.8% TAE agarose gel to visualize their cleavage efficiency for SpCas9. AgeI (catalog no. R0552S; New England Biolabs) digested pX551 plasmid was used as a positive control. Four SpCas9 sgRNAs were then cloned into a pX552-CMV-GFP plasmid (modified from Addgene #60958, by replacing the hSyn promoter with a CMV promoter) at the SapI restriction site for *in vitro* validation. Subsequently, the selected SpCas9 sgRNA (SpCas9 sgRNA4) was sub-cloned into a AAV package plasmid (pX552-hsyn-mCherry-YFP sgRNA2, sgRNA6 or pX552-LacZ sgRNA) at the MluI (catalog no. R3198; New England Biolabs) restriction site to generate YFP or LacZ targeting kamikaze CRISPR/Cas construct. For *in vitro* validation, pX551-CMV-SpCas9 plasmid was modified from pX551 plasmid by replacing the MeCP2 promoter with a CMV promoter.

### Cell culture and transfection

Stable *YFP* expressing HEK293A cells were generated using a lentivirus as previously described.^10,15^ HEK293A-YFP cells were maintained in Dulbecco's modified Eagle's media (DMEM) (catalog no. 11965118; Life Technologies Australia, Scoresby, VIC, Australia) supplemented with 10% fetal calf serum (Sigma-Aldrich, St. Louis, MO, USA), 2 mM glutamine (catalog no. 2503008; Life Technologies Australia), 50 U/mL penicillin-streptomycin (catalog no. 15070063; Life Technologies Australia) in a humidified 5% CO_2_ atmosphere at 37 °C. Transfection was undertaken with FuGENE-HD transfection reagent (catalog no. E2311; Promega Australia, Alexandria, NSW, Australia) according to the manufacturer’s instruction. Briefly, HEK293A-YFP cells were seeded onto a 6-well plate (2.5×10^5^ per well) 24 hours before transfection. A mixture of 7.5 μL FuGENE-HD transfection reagent with 1500 ng plasmid in 150 μL Opti-MEM (catalog no. 11058021; Life Technologies Australia) was added into each well. For *in vitro* validation of SpCas9 sgRNA, cells were collected for western blot analysis at day 3 after transfection; for *in vitro* time course analysis, cells were harvested at day 1, 2, 3 and 5 after transfection.

### Western blot analysis

Cells were collected and lysed in ice-cold Cell Lysis Buffer (Catalog no.89900; Thermo Scientific, Waltham, MA, USA) and sonicated for 10 seconds by an ultrasonic cell disruptor (MISONIX Microson XL 2000; Qsonica, Newtown, CT, USA). Total protein was quantified by a Bio-Rad protein assay (Catalog no. 5000006; BIO-RAD, Hercules, CA, USA) using microplate reader (Infinite M1000 Pro; TECAN, Männedorf, Switzerland). A total of 10 μg protein samples were separated using NuPAGE™ Novex™ 4-12% Bis-Tris Protein Gels (catalog no. NP0321BOX; Life Technologies Australia) and transferred to polyvinylidene fluoride membranes (catalog no. 162-0177; BIO-RAD) using the XCell II™ Blot Module (Life Technologies Australia). Membranes were blocked with 5% skim milk in TBS-T (10 mM Tris, 150 mM NaCl, and 0.05% Tween-20) at room temperature for 1 hour and then incubated with mouse monoclonal SpCas9 antibody (1:1000 dilution; MAC133, lot number 2591899; Millipore, Billerica, MA, USA) or mouse monoclonal β-actin antibody (1:2000 dilution; MAB 1501, lot number 2722855; Millipore) at room temperature for 1 hour. Membranes were washed, further incubated with horseradish peroxidase-conjugated goat anti-mouse secondary antibody (1:5000 dilution; catalog no. A-11045; Life Technologies Australia) at room temperature for 1 hour, and developed using the Amersham ECL Prime Western Blotting Detection kit (catalog no. RPN2232; GE Healthcare Australia, Parramatta, NSW, Australia). The relative levels of SpCas9 protein of each sample was quantified using densitometry analysis (ImageJ software-gels analysis) with normalization to β-actin.

### YFP detection

YFP expressing HEK293A cells were trypsinized and harvested in PBS. The percentage of YFP positive cells was analyzed with a MACSQuant Flow Cytometer (Miltenyi Biotec Australia, Macquarie Park, NSW, Australia), and data were analyzed using cell cycle analysis software (FlowJo^®^; FlowJo LLC, Ashland, OR, USA).

### AAV production

Recombinant AAV2 viruses were produced in HEK293D cells (kindly provided by Ian Alexander, Children's Medical Research Institute, Westmead, NSW, Australia) packaging either pX551 plasmid, containing SpCas9, or pX552-mCherry plasmid with the respective sgRNAs (spCas9, YFP or LacZ sgRNA), pseudoserotyped with the AAV2 capsid (pXX2), and purified using a AAV2pro Purification Kit (catalog no. 6232; Clontech Laboratories, Mountain View, CA, USA) as previously described.^10,15,16^ Viral titer was determined by real-time quantitative PCR using a Fast SYBR Green Master Mix (catalog no. 4385612; Life Technologies Australia) with the pX551 or pX552 forward and reverse primers (Supplementary Table 1).

### Intravitreal Injection

For our *in vivo* time course analysis, a total of 76 C57BL/6 adult mice, aged between 12 and 14 weeks were randomly separated into two groups, to receive either AAV2-SpCas9+AAV2-YFP sgRNA2 (n= 39) or AAV2-SpCas9+AAV2-SpCas9 sgRNA/YFP sgRNA2 (n= 37). For the YFP disruption experiments, a total of 49 *Thy1-YFP* transgenic mice, aged between 16 and 20 weeks, were randomly allocated into three groups; those receiving AAV2-SpCas9+AAV2-YFP sgRNA2 (n= 17), AAV2-SpCas9+AAV2-SpCas9 sgRNA/YFP sgRNA2 (n=17) or AAV2-SpCas9+AAV2-SpCas9 sgRNA/LacZ sgRNA (n= 15). In addition, another 29 *Thy1-YFP* transgenic mice were used to test different YFP sgRNAs. These mice were randomly allocated into three groups; those receiving AAV2-SpCas9+AAV2-YFP sgRNA6 (n= 9), AAV2-SpCas9+AAV2-SpCas9 sgRNA/YFP sgRNA6 (n= 10) or AAV2-SpCas9+AAV2-SpCas9 sgRNA/LacZ sgRNA (n= 10).

Mice were anesthetized by intraperitoneal injection of ketamine (60 mg/kg) and xylazine (10 mg/kg).^15^ Intravitreal injection was performed under a surgical microscope. After a small puncture was made through the conjunctiva and sclera using a 30-gauge needle, a hand-pulled glass micropipette connected to a 10 μL Hamilton syringe (Bio-Strategy, Broadmeadows, VIC, Australia) was inserted into the vitreous. A total of 1 μL dual-viral suspension (AAV2-SpCas9: 2.5×10^9^ vector genomes vg/μL with AAV2-YFP sgRNA: 2.5×10^9^ vg/μL, AAV2-SpCas9 sgRNA/YFP-sgRNA: 2.5×10^9^ vg/μL or AAV2-SpCas9 sgRNA/LacZ sgRNA: 2.5×10^9^ vg/μL) was injected into one eye of each mouse using a UMP3-2 Ultra Micro Pump (World Precision Instruments, Sarasota, FL, USA) at a rate of 200 nL/s. Any issues with the injection, including backflow upon removal of the needle, hemorrhaging of the external or internal vessels, retinal detachment were recorded and eyes were excluded from the study.

### Electroretinography (ERG) and Optical Coherence Tomography (OCT)

At 8 weeks following injection, mice underwent overnight dark-adaptation (~12 hours), followed by electroretinography assessment under fully dark-adapted conditions. Details for functional assessment have been outlined previously,^17^ with the exception that the reference chloride silver electrode was placed around the outside of the eye. ERG analysis was as previously described^17^ and returned the photoreceptor (a-wave), bipolar cell (b-wave), and ganglion cell dominated (scotopic threshold response, STR) components of the waveform. Group data are given as mean (± standard error of the mean).

Following ERG recordings, retinal images were obtained using a spectral domain-OCT (Bioptigen, Inc., Morrisville, NC, USA). Mice were positioned to capture Optic Nerve Head (ONH) centred 1.4 mm-wide horizontal B-scans (consisting of 1000 A-scans). ImageJ software (https://imagej.nih.gov/ij/) was used in a masked fashion to quantify total retinal thickness (from the inner limiting to Bruch’s membrane), retinal nerve fibre layer thickness (from the inner limiting membrane to the inner aspect of the inner plexiform layer) and outer retinal thickness (from Bruch’s membrane to the outer plexiform layer) in each eye.

### Retinal flat-mount, imaging and counting

Eyes were removed, fixed in ice-cold 4% paraformaldehyde for 1 hour and dissected under a dissecting microscope. After removing the cornea, iris and lens, four equally spaced radial relaxing incisions, extending two thirds of the way from the retinal periphery to the ONH, were made. The sclera and choroid were then removed along with residual vitreous and hyaloid vessels, leaving only the retina. The fully dissected retina was stained with NucBlue™ Live ReadyProbes™ Reagent (catalog no. R37605; Life Technologies Australia) as a nuclear counterstain. Retinal images were captured by a fluorescence microscope (Zeiss Axio Imager Microscope; Carl-Zeiss-Strasse, Oberkochen, Germany) equipped with a charge-coupled digital camera (Axiocam MRm, Zeiss) and image acquisition software (ZEN2, Zeiss) as previously described.^10^

The efficiency of YFP disruption was quantified using individual fluorescent images captured at ×400 magnification. A total of 24 images from three flat-mounted eyes treated with SpCas9 sgRNA/LacZ sgRNA, 36 images from five flat-mounted eyes treated with SpCas9 sgRNA/YFP sgRNA2 and 36 images from five flat-mounted eyes treated with YFP sgRNA2 were quantified manually using ImageJ v1.49 by an experienced grader (FL), masked to treatment status. For the second experiment with YFP sgRNA6, 16 images from three flat-mounted eyes treated with SpCas9 sgRNA/LacZ sgRNA, 38 images from five flat-mounted eyes treated with SpCas9 sgRNA/YFP sgRNA6 and 36 images from six flat-mounted eyes treated with YFP sgRNA6 were quantified. Efficiency for each treatment group was determined as the proportion of YFP-negative cells relative to mCherry-expressing cells as previously described.^10^

### Quantitative PCR

Total RNA from mouse retinas were extracted and purified using commercial kits (RNeasy Mini Kit; catalog no. 74104; Qiagen, Chadstone, VIC, Australia) in accordance with the manufacturer's instructions. RNA was subsequently reverse-transcribed into complementary DNA (cDNA) using a high-capacity RT kit (catalog no. 4374996; Life Technologies Australia) and quantitative PCR was performed using a Fast SYBR Green Master Mix (catalog no. 4385612; Life Technologies Australia) with the SpCas9 forward and reverse primers as well as mCherry forward and reverse primers (Supplementary Table 1). The relative expression levels of SpCas9 was calculated using the ^∆∆^Ct method with normalization to mCherry.

### Statistical Analysis

All statistical analyses were performed using Prism 7 software (GraphPad Software, Inc., La Jolla, CA, USA). Group data are represented as mean ± SEM unless otherwise noted. Mean data were analyzed with unpaired t-tests, one-way or two-way analysis of variance (ANOVA) followed by post-hoc analysis (GraphPad Prism 7.0). A value of p < 0.05 was taken to be statistically significant.

## RESULTS

### Generation and validation of kamikaze CRISPR/Cas construct *in vitro*

We first validated four sgRNAs (Figure 2A) for SpCas9 targeting using an *in situ* cleavage assay. Robust cleavage of the SpCas9 plasmid (pX551) was found when each of the four designed SpCas9 sgRNAs were introduced to recombinant SpCas9 protein (Figure 2B). We further confirmed the efficacy of SpCas9 gene perturbations by transfection of the SpCas9 expression construct (pX551-CMV-SpCas9) together with SpCas9 targeting CRISPR/Cas constructs carrying different SpCas9 sgRNA (pX552-SpCas9 sgRNA1-4) in HEK293A cells. SpCas9 sgRNA4 had a clear destructive effect on SpCas9 (Figure 2C), reduction of SpCas9 protein, as well as having a lower off-target score against the human genome as predicted by a web-based CRISPR design program (http://crispr.mit.edu). A time course analysis showed that SpCas9 protein was progressively reduced (46% at day 1, 77% at day 2 and 86% at day 3, 56% at day 5) in cells following the transfection of selected SpCas9 targeting CRISPR/Cas construct (SpCas9 sgRNA4) compared to LacZ sgRNA control (Figures 2D and 2E).

**Figure 2.**
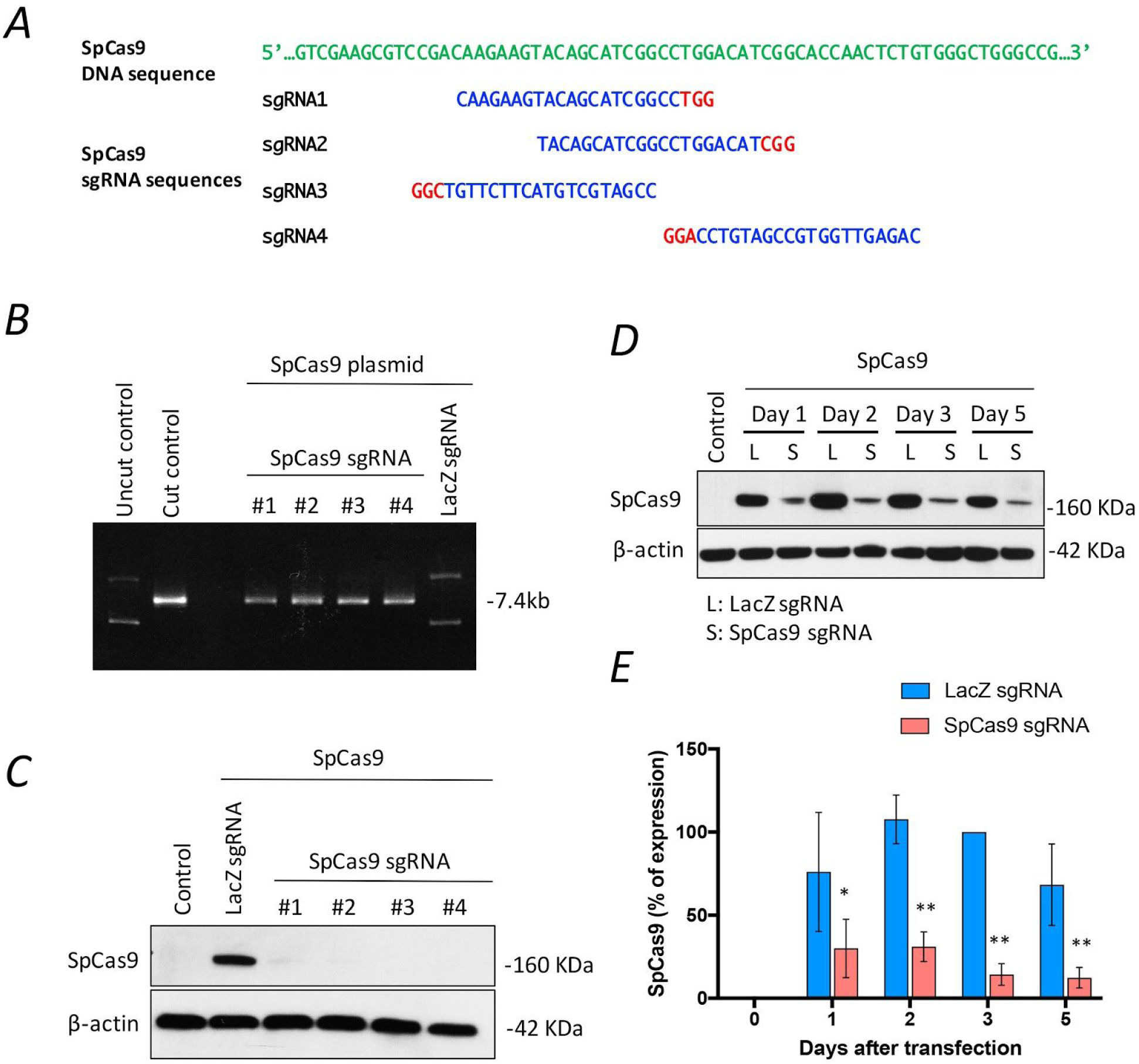
Design and validation of SpCas9 sgRNA. (A) Schematic diagram of SpCas9 sgRNA design. Green: SpCas9 sequence. Blue: selected SpCas9 sgRNA targeted sites. Red: PAM sequences. (B) *In situ* validation of SpCas9 sgRNAs. (C) *In vitro* validation of SpCas9 sgRNAs. Representative western blot of SpCas9 protein expression in cells co-transfected with SpCas9 and the individual SpCas9 sgRNA plasmids. (D) Representative western blot of the time course of SpCas9 expression. Cells were harvested on day 1, 2, 3 and 5 after transfection with SpCas9 and selected SpCas9 sgRNA (#4) plasmids. (E) Relative fold change of SpCas9 expression normalized to β-actin, as a function of days treatment with SpCas9 sgRNA or LacZ sgRNA. Mean ± SEM for 3 independent replicates. Statistical analysis between groups was performed using two-way ANOVA followed by Sidak's multiple comparisons test (*p < 0.05, **p < 0.001).

Next we re-engineered our Kamikaze CRISPR/Cas construct with YFP targetting sgRNAs or a LacZ targeting sgRNA (Figure 3A), and the efficacy of YFP gene disruption in the YFP-expressing HEK293A cells was assessed. We observed a significant reduction of SpCas9 protein in cells that had received the kamikaze CRISPR/Cas construct compared to those cells that had received conventional CRISPR/Cas construct 2 days after transfection (Figure 3B). In terms of efficiency, our result indicated that the percentage of YFP-expressing cells was significantly reduced in cells transfected with the YFP targeting kamikaze CRISPR/Cas constructs (YFP sgRNA2: 7.2±0.6% and YFP sgRNA6: 6.5±3.2% respectively), compared to LacZ targeting kamikaze (97.9±1.2%) or LacZ targeting (95.8±0.4%) CRISPR/Cas construct 10 days after transfection (Figure 3C). Similarly, a lower percentage of YFP expressed cells could also be found in cells transfected with the YFP targeting CRISPR/Cas construct (YFP sgRNA2: 11.9±4.7% and YFP sgRNA6: 4.7±1.8% respectively; Figure 3C).

**Figure 3.**
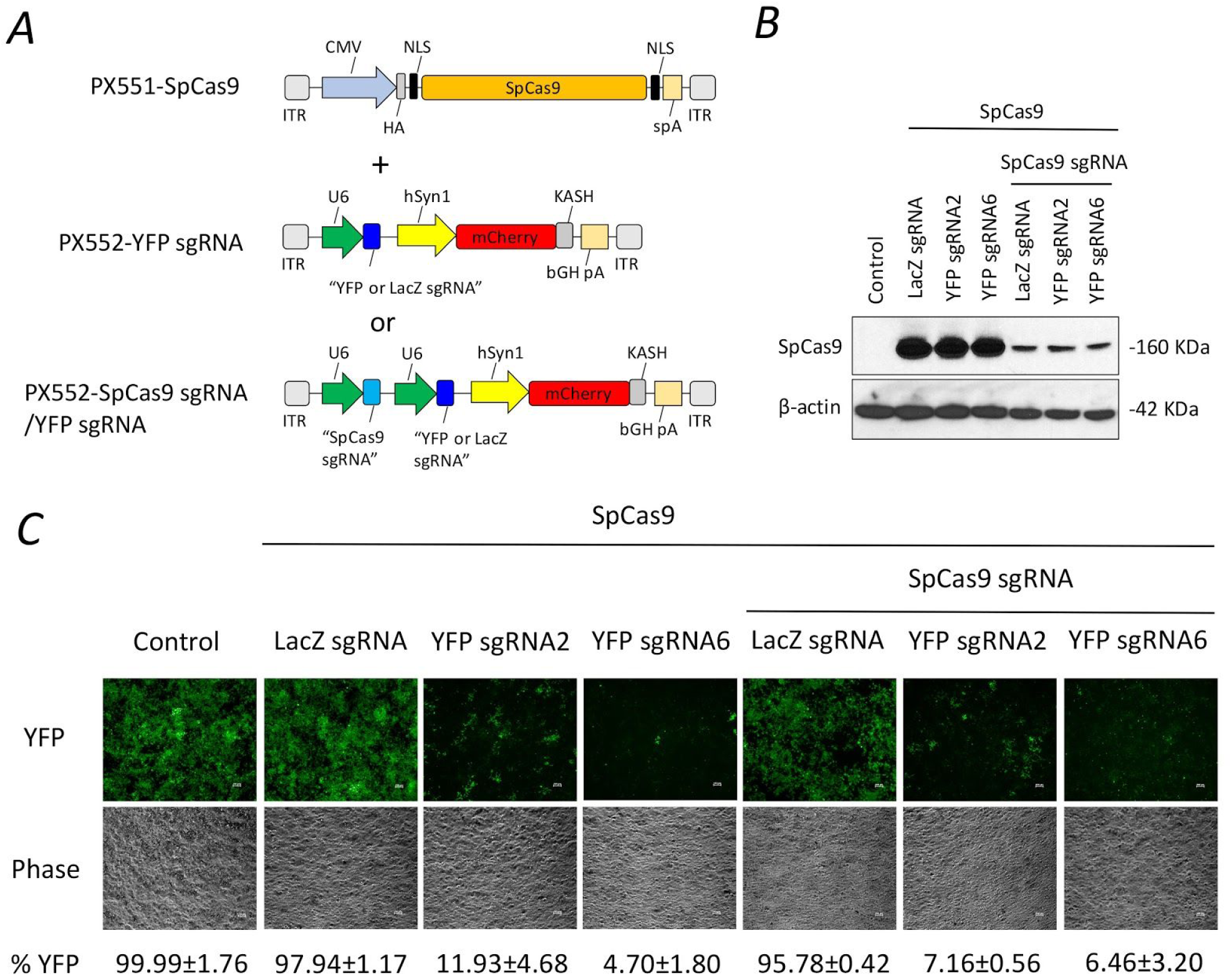
*In vitro* validation of kamikaze CRISPR/Cas construct. (A) Schematic of plasmid constructs for *in vitro* validation. (B) Representative Western blots of SpCas9 protein expression in cells co-transfected with SpCas9 and kamikaze (SpCas9 sgRNA/YFP sgRNA and SpCas9 sgRNA/LacZ sgRNA) or non-kamikaze (YFP sgRNA and LacZ sgRNA) constructs. (C) Representative images of YFP expression in cells co-transfected with kamikaze (SpCas9 sgRNA/YFP sgRNA and SpCas9 sgRNA/LacZ sgRNA) or non-kamikaze (YFP sgRNA and LacZ sgRNA) constructs. Percentage YFP disruption was assessed by FACS at 10 day after transfection. scale bar: 100 μm. Mean ± SEM for 2 independent replicates.

### *In vivo* delivery of kamikaze CRISPR/Cas construct in the mouse retina

To evaluate whether the reduction of SpCas9 expression by the kamikaze CRISPR/Cas construct compromises on-target editing efficiency *Thy1-YFP* mice received a single intravitreal injection of a dual viral suspension of AAV2-SpCas9 along with the YFP targeting kamikaze CRISPR/Cas construct (AAV2-SpCas9 sgRNA/YFP sgRNA) or the LacZ targeting kamikaze CRISPR/Cas construct (AAV2-SpCas9 sgRNA/LacZ sgRNA) or a single YFP targeting CRISPR/Cas construct as a positive control (AAV2-YFP sgRNA2). Eight weeks following treatment, images from the retinal flat-mounts showed that there were fewer YFP-positive cells among mCherry positive cells in mice that had received either AAV2-SpCas9 sgRNA/YFP sgRNA or AAV2-YFP sgRNA2 compared to AAV2-SpCas9 sgRNA/LacZ sgRNA (Figure 4A). Specifically, the proportion of retinal YFP/mCherry-expressing cells was reduced to 5.5±1.4% in AAV2-SpCas9 sgRNA/YFP sgRNA2-treated retina and 7.3±1.3% in AAV2-YFP sgRNA2-treated retina, compared with 38.2±1.7% in AAV2-SpCas9 sgRNA/LacZ sgRNA treated eyes. Overall, there was a 85.5% (95% CI: 78.4-92.6) and 80.9% (95% CI: 74.3-87.5) reduction in YFP positive cells in AAV2-SpCas9 sgRNA/YFP sgRNA- and AAV2-YFP sgRNA2-treated retinas, respectively, compared to tAAV2-SpCas9 sgRNA/LacZ sgRNA-treated eyes (Figure 4B). No significant difference in the percentage of YFP disruption was found in between AAV2-YFP sgRNA2- and AAV2-SpCas9 sgRNA/YFP sgRNA2-treated retinas (P=0.62; Figure 4B). This was confirmed by using an alternate YFP targeting sgRNA (YFP sgRNA6), where the proportion of retinal YFP/mCherry-expressing cells was 17.0±1.3% in AAV2-SpCas9 sgRNA/YFP sgRNA6-treated retina and 20.6±1.2% in AAV2-YFP sgRNA6-treated retina, compared with 40.8±2.0% AAV2-SpCas9 sgRNA/LacZ sgRNA-treated eyes. This represents a relative reduction of 49.5% (95% CI: 43.5-55.5) and 58.3% (95% CI: 56.4-62.0) in AAV2-SpCas9 sgRNA/YFP sgRNA6- and AAV2-YFP sgRNA6-treated retinas compared to those that had received AAV2-SpCas9 sgRNA/LacZ sgRNA, respectively (Supplementary Figure 1).

**Figure 4.**
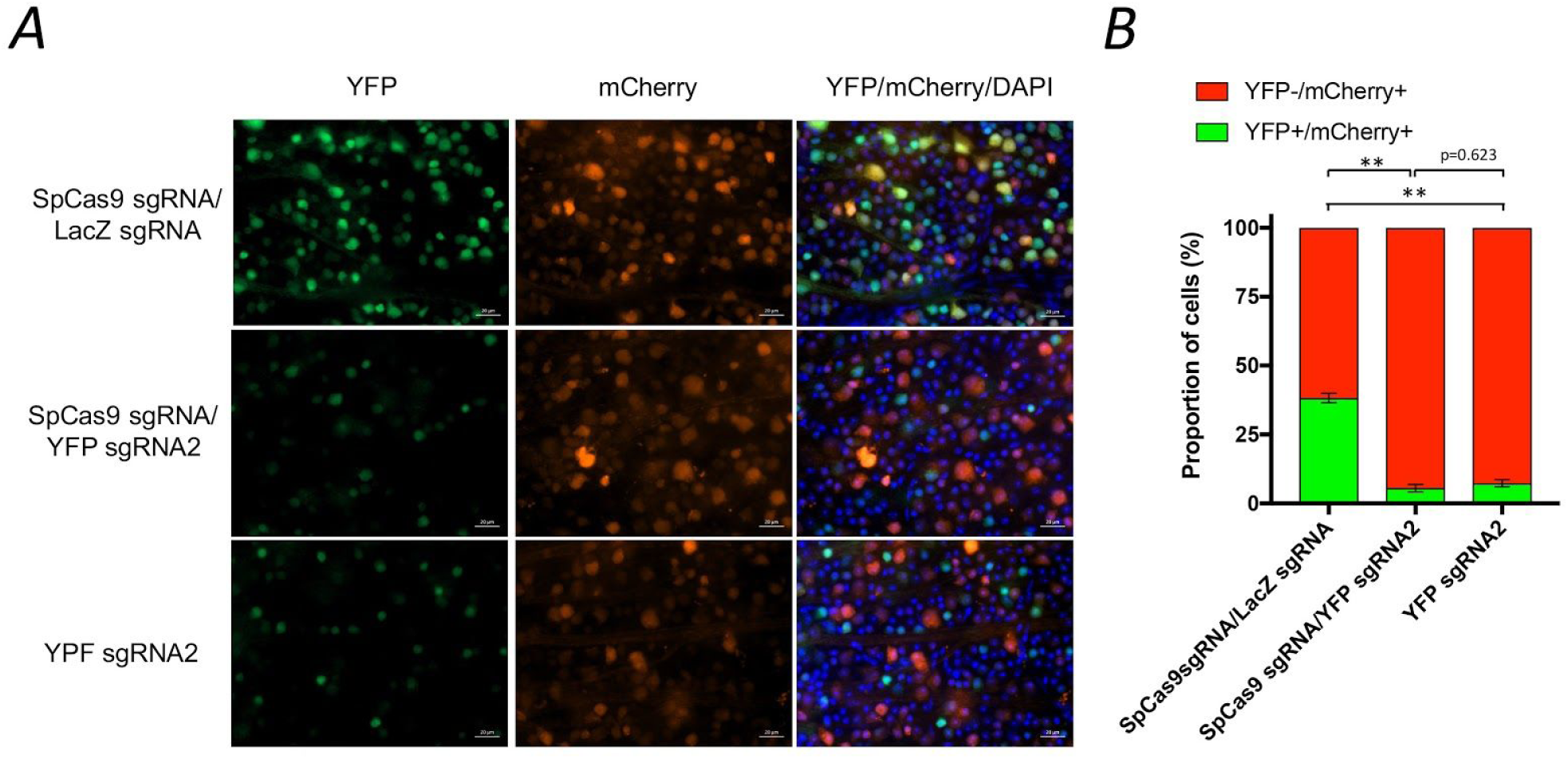
Kamikaze CRISPR/Cas-mediated genome editing of retinal cells *in vivo*. (A) High magnification of retinal flat-mount images, showing differences in YFP expression following AAV2-mediated delivery of SpCas9 sgRNA/YFP sgRNA (n= 5), YFP sgRNA2 (n= 5) or SpCas9 sgRNA/LacZ sgRNA (n= 3). scale bar: 20 μm. (B) Percentage YFP disruption was assessed by manual cell counting. Mean ± SEM for 3-5 independent replicates. Statistical analysis between groups was performed using one-way ANOVA followed by Tukey's multiple comparisons test (**p < 0.001).

### Retinal function and structure assessment by electroretinography (ERG) and optical coherence tomography (OCT)

To evaluate the effect of our “kamikaze” CRISPR/Cas construct on retinal function and structure, ERG and OCT were performed at 8 weeks after intravitreal injection of viral suspensions in *Thy1-YFP* mouse. Group averaged waveforms elicited using bright and dim flashes of light along with the group averaged data from eye injected with YFP targeting kamikaze-CRISPR/Cas constructs (AAV2-SpCas9 sgRNA/YFP sgRNA2, Figures 5A and 5B) and YFP targeting CRISPR/Cas constructs (AAV2-YFP sgRNA2, Figures 5E and 5F) suggest that both treatments affected retinal function when compared with the contralateral control eyes (Figure 5A-F and Supplementary Figures 2-4). LacZ targeting kamikaze-CRISPR/Cas construct (AAV2-SpCas9 sgRNA/LacZ sgRNA) treated eyes retained normal retinal function (Figure 5C and 5D). OCT analysis suggest that none of the CRISPR/Cas constructs negatively impacted retinal structure, as there as no significant differences in retinal nerve fibre layer and total retinal thickness between the vehicle and viral-injected eyes of all three groups (Figure 5G-I).

**Figure 5.**
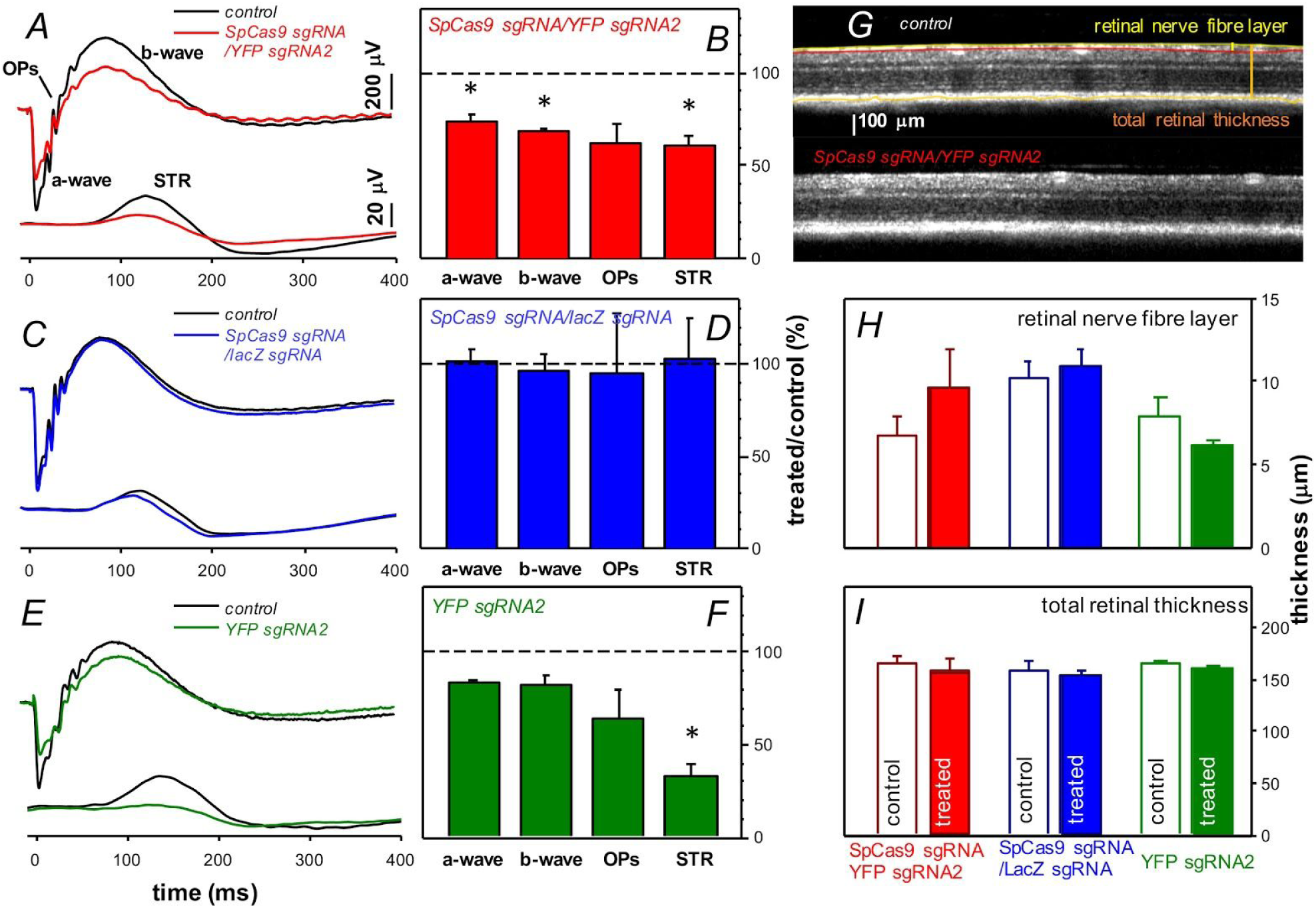
Long-term effect of AAV2-mediated CRISPR/Cas administration on retinal function. Averaged ERG waveforms at selected intensities for control (black traces) and SpCas9 sgRNA/YFP sgRNA2 (n= 4, red traces; A), SpCas9 sgRNA/LacZ sgRNA (n = 4, blue traces; C) and YFP sgRNA2 (n= 5, green traces; E) injected eyes. Group average (± SEM) photoreceptoral (a-wave), bipolar cell (b-wave), amacrine cell (oscillatory potentials, OPs) and ganglion cell (scotopic threshold response, STR) amplitude relative to contralateral control eyes (%) for each group (B, D and F). Effect of SpCas9 sgRNA/YFP sgRNA2, SpCas9 sgRNA/LacZ sgRNA and YFP sgRNA2 on retinal structure measured with OCT (G). Group average (± SEM) retinal nerve fibre layer thickness (H) for SpCas9 sgRNA/YFP sgRNA2 treated (filled red, n= 4) and their contralateral controls (unfilled red, n= 4), SpCas9 sgRNA/LacZ sgRNA treated (filled blue, n= 4) and their contralateral controls (unfilled blue, n= 4) and YFP sgRNA2 treated (filled green, n= 5) and their contralateral controls (unfilled green, n=5). Total retinal thickness (I). Statistical analysis between injected and control eyes was performed using two-tailed Student t-test (*p<0.05).

## DISCUSSION

This study builds on our recent work using AAV2-mediated CRISPR/Cas to edit genes in mouse retina. While CRISPR/Cas9-mediated genome editing has shown promise for correcting disease-causing mutations, the potential for genotoxic effects with prolonged expression of CRISPR/Cas9 poses a significant barrier to the clinical utility of this technology. A recent study suggest that there can be an unexpectedly high number of single-nucleotide variants in mice that had undergone CRISPR/Cas9-mediated genome editing.^18^ Several strategies have been employed in an attempt to avoid off-target cleavage, including improved guide RNA design,^19,20^ or modification of Cas9 enzymes.^21,22^ Such approaches do not avoid accumulation of Cas9, which can increase the overall chance of off-target cleavage. Our approach was to employ a self-destructive kamikaze CRISPR/Cas system that disrupts the CRISPR/Cas enzyme itself after the active protein has been expressed. Unlike other approaches, most of which act to control the activity of the CRISPR/Cas system via chemical,^23,24^ and biophysical^25,26^ modulation of Cas9, our kamikaze CRISPR/Cas system can significantly reduce accumulation of Cas9, and thus off-target cleavage, without dramatically compromising the efficiency of on-target editing. This approach is similar to that used by Merienne and colleagues,^27^ who demonstrated that progressively inactivating the nuclease using a Cas9 self-inactivating editing system resulted in a lower frequency of off-target cleavage in human iPSCs-derived neurons *in vitro* and in mouse brains *in vivo*^21^ This highlights the potential for a viral-mediated self-destructive CRISPR/Cas systems to be potentially used as a safer tool for *in vivo* genome editing.

We observed a reduction in the efficiency of SpCas9 gene perturbations with the kamikaze CRISPR/Cas system between *in vitro* (85.7%) and *in vivo* (63.8% at 8 weeks after injection; supplementary Figure 5) models. This difference may be due to the different promoters and delivery systems used *in vitro* and *in vivo*. A constitutively ubiquitous cytomegalovirus (CMV) promoter was used to drive the expression of SpCas9 targeting sgRNA in the *in vitro* experiment, whereas the MeCP2 promoter was used to achieve specific neuronal expression *in vivo*. Lower *in vivo* efficiency may also explain why levels of SpCas9 protein in the mouse retina were undetectable by western blot analysis (Supplementary Figure 6). Another difference was that a dual AAV2 vector system was employed to deliver the kamikaze CRISPR/Cas construct *in vivo*. We and others have recently demonstrated that CRISPR/Cas9 delivered using a dual AAV2 vector can effectively edit the genome in a number of organs in adult mice.^10,28-30^ However, expression of the CRISPR/Cas9 machinery requires the receipt of both Cas9 and sgRNA expression cassettes from two separate viral vectors, which may significantly reduce editing efficiency.^28^ Although a single viral vector system employing Cas9 orthologs such as SaCas9^31^ or CjCas9^32^ may provide better *in vivo* editing efficiency, dual-vector systems may still be required for mutation correction as they enable delivery of donor templates and appropriate promoter elements.

An unexpected reduction in retinal function was observed 8 weeks after injection of AAV2-SpCas9 sgRNA/YFP sgRNA2 or AAV2-YFP sgRNA2. Interestingly, retinal function was unaffected in mice treated with AAV2-SpCas9 sgRNA/LacZ sgRNA, therefore, deficits in retinal function may be related to the YFP targeting sgRNA. To explore this possibility a further *in vivo* study was undertaken, employing a different YFP sgRNA (sgRNA6), which targets another region of the YFP sequence. Data shown in Supplementary Figures 7-9, shows that a significant decrease in retinal function was still present in AAV2-SpCas9 sgRNA/YFP sgRNA6 and AAV2-YFP sgRNA6 treated mice. These results indicate that potential retinal dysfunction was unlikely to have resulted from off-target effects of YFP sgRNA (either 2 or 6). However, without further confirmation by whole-genome sequencing on these mice, we cannot completely rule out off-target effects from YFP sgRNA.

In addition to potential off-target effects of YFP sgRNA, accumulation of non-functional fluorescent proteins resulting from CRISPR/Cas9 editing may be possible. Although fluorescence proteins such as GFP and YFP have been widely used in neuroscience research,^33^ accumulation of non-functional proteins resulting from on-target deletions (indel) may have a deleterious effect on retinal protein homeostasis.^34^ Moreover, a recent study also indicated that large on-target deletions could lead to potential genotoxicity.^29^ Whether such mechanisms account for the functional deficits observed in our study requires further investigation. Nevertheless these data highlight the need for careful design of AAV-CRISPR/Cas9 system for application in complex tissues.

In summary, we describe and characterise a self-destructive “kamikaze” CRISPR/Cas system for *in vivo* genome editing in the retina. This self-destructive kamikaze CRISPR/Cas system can effectively reduce the expression of SpCas9 in the mouse retina, without substantially sacrificing on-target editing efficiency. Therefore, our AAV2-mediated self-destructive CRISPR/Cas may be a robust tool for genome editing in the retina, especially when combined with high fidelity forms of CRISPR/Cas.

## ACKNOWLEDGMENTS

This work was supported by funding from a Bayer Global Ophthalmology Award, the Ophthalmic Research Institute of Australia, an Australian National Health and Medical Research Council (NHMRC) grant (APP1123329), an NHMRC Practitioner Fellowship (AWH, 1103329), and an Australian Research Council Future Fellowship (AP, FT140100047). CERA receives Operational Infrastructure Support from the Victorian Government.

## SUPPLEMENTARY METHODS AND RESULTS

**Supplementary Table S1.**
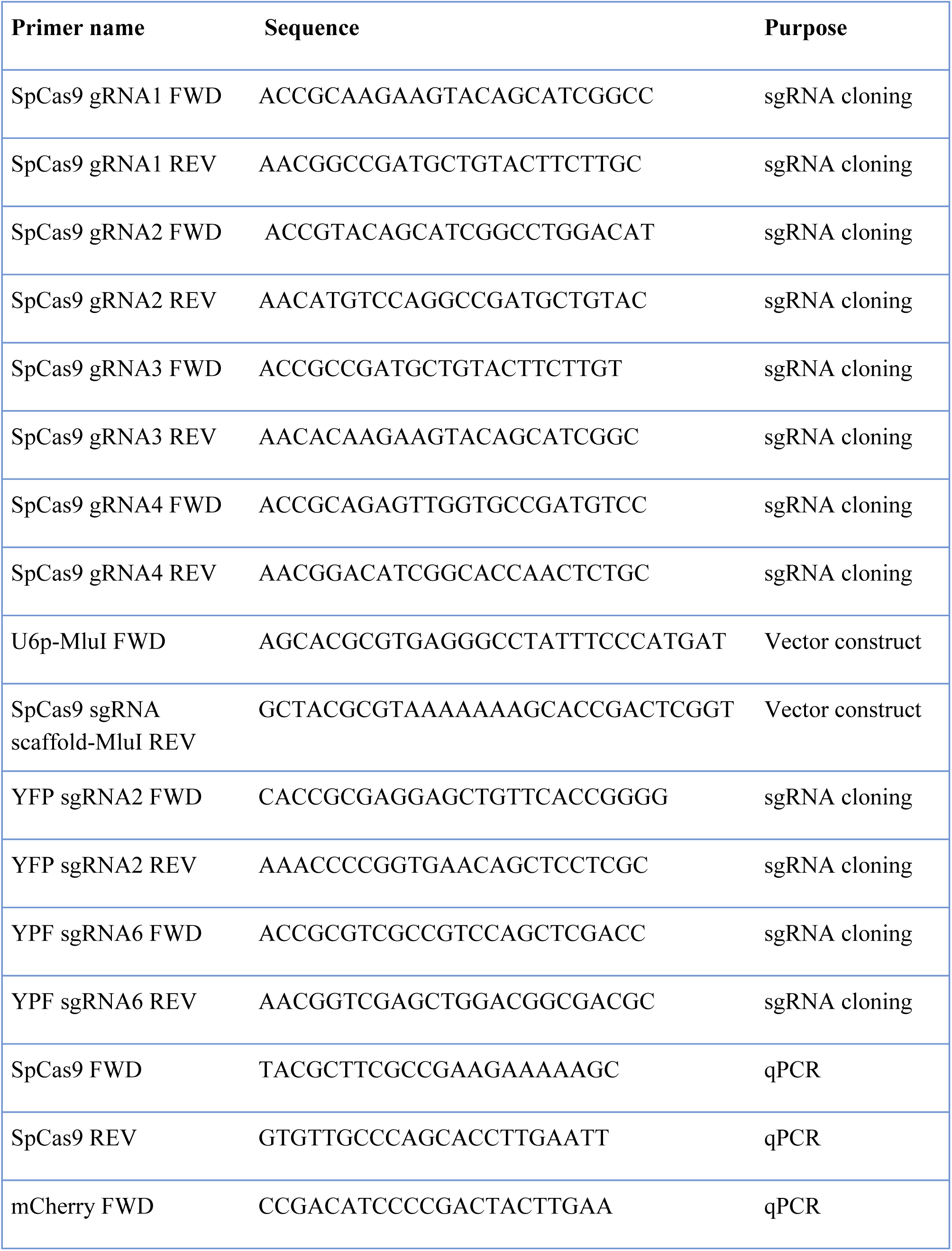
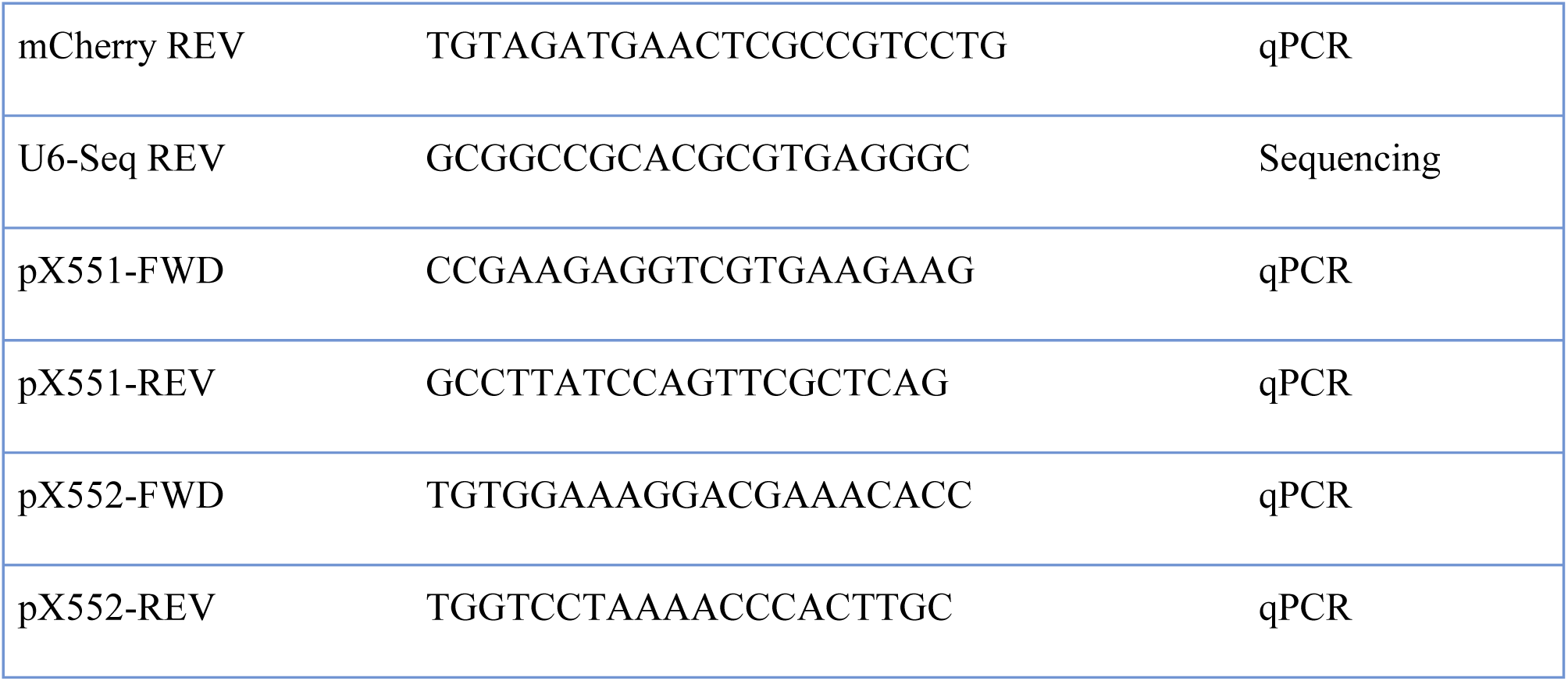
Sequence of primers for sgRNA cloning, vector construction, sequencing and qPCR analysis.

**Supplementary Figure 1.**
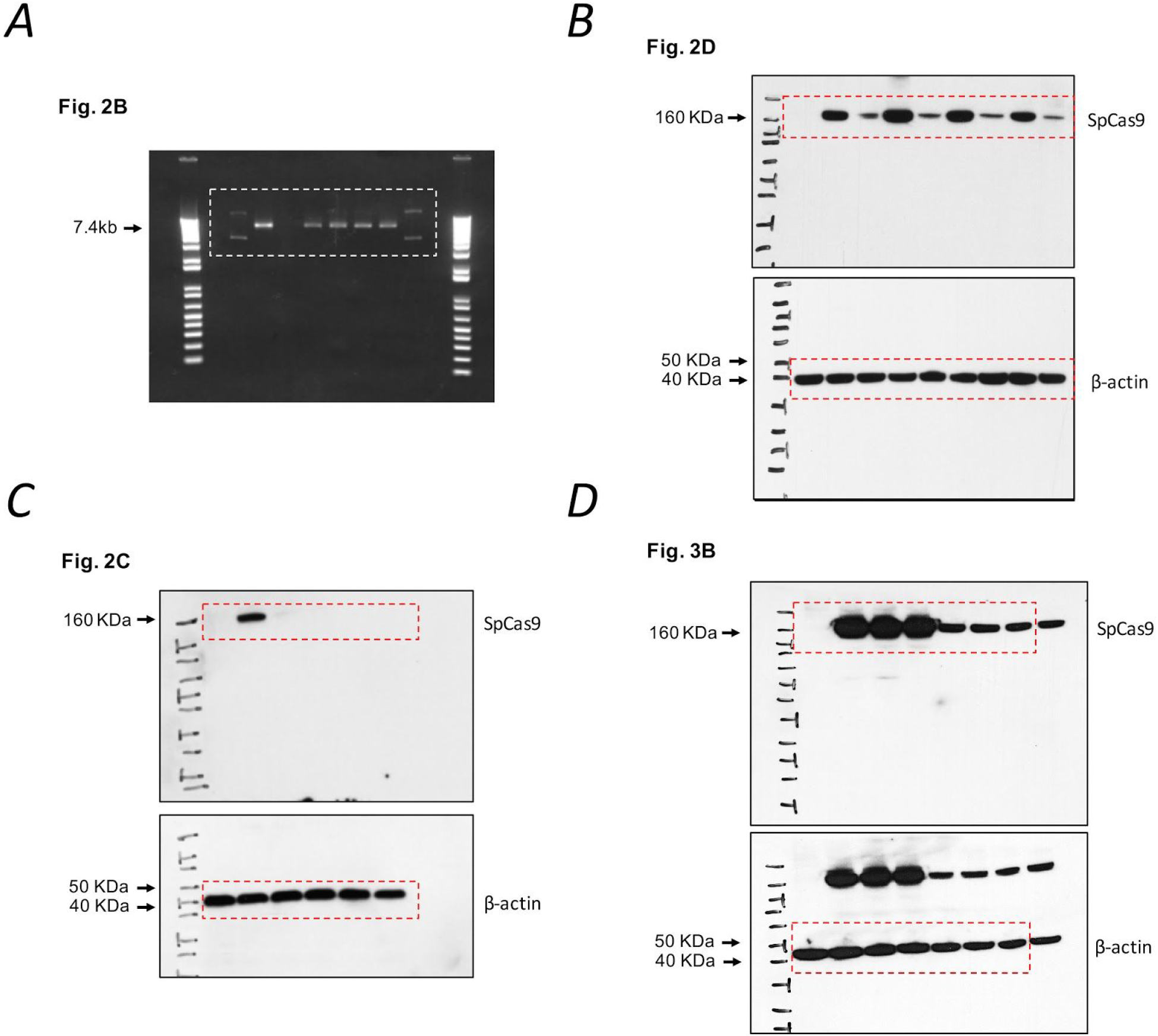
Uncropped agarose gel and western blot images. (A) *In situ* test of designed SpCas9 sgRNAs. (B) *in vitro* validation of designed SpCas9 sgRNAs. (C) Time course of SpCas9 expression. **(D)** *in vitro* validation of YFP targeting kamikaze CRISPR constructs. The β-actin membrane was reprobing without stripping.

**Supplementary Figure 2.**
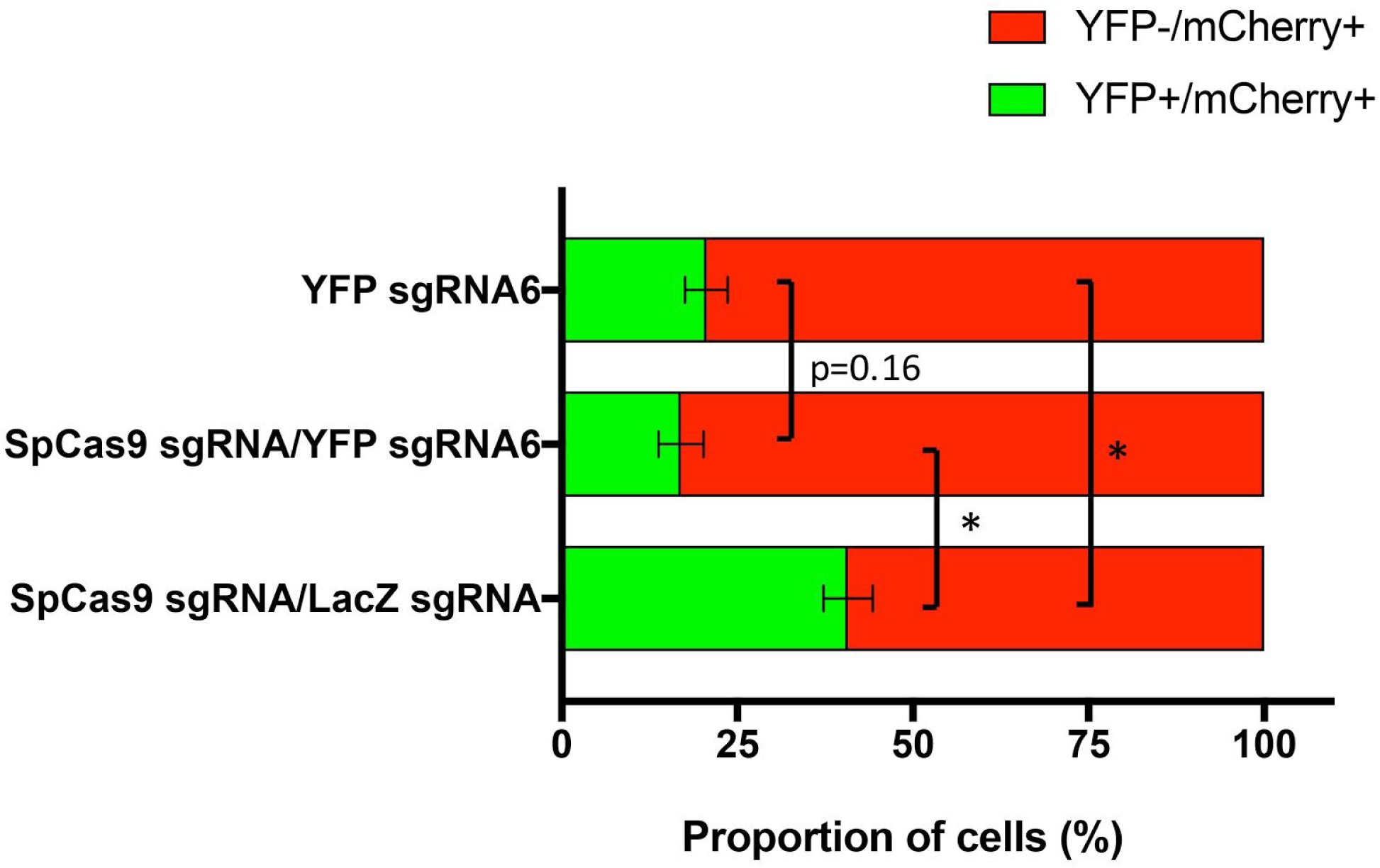
Quantification of YFP disruption in the retina. The percentage of YFP disruption following AAV2-mediated delivery of SpCas9 sgRNA/YFP sgRNA6, YFP sgRNA6 or SpCas9 sgRNA/LacZ sgRNA was assessed by manual cell counting. Representative data are shown for 3-5 retinas and expressed as the Mean ± SEM. Statistical analysis between groups was performed using one-way ANOVA followed by Tukey's multiple comparisons test (*p < 0.05).

**Supplementary Figure 3.**
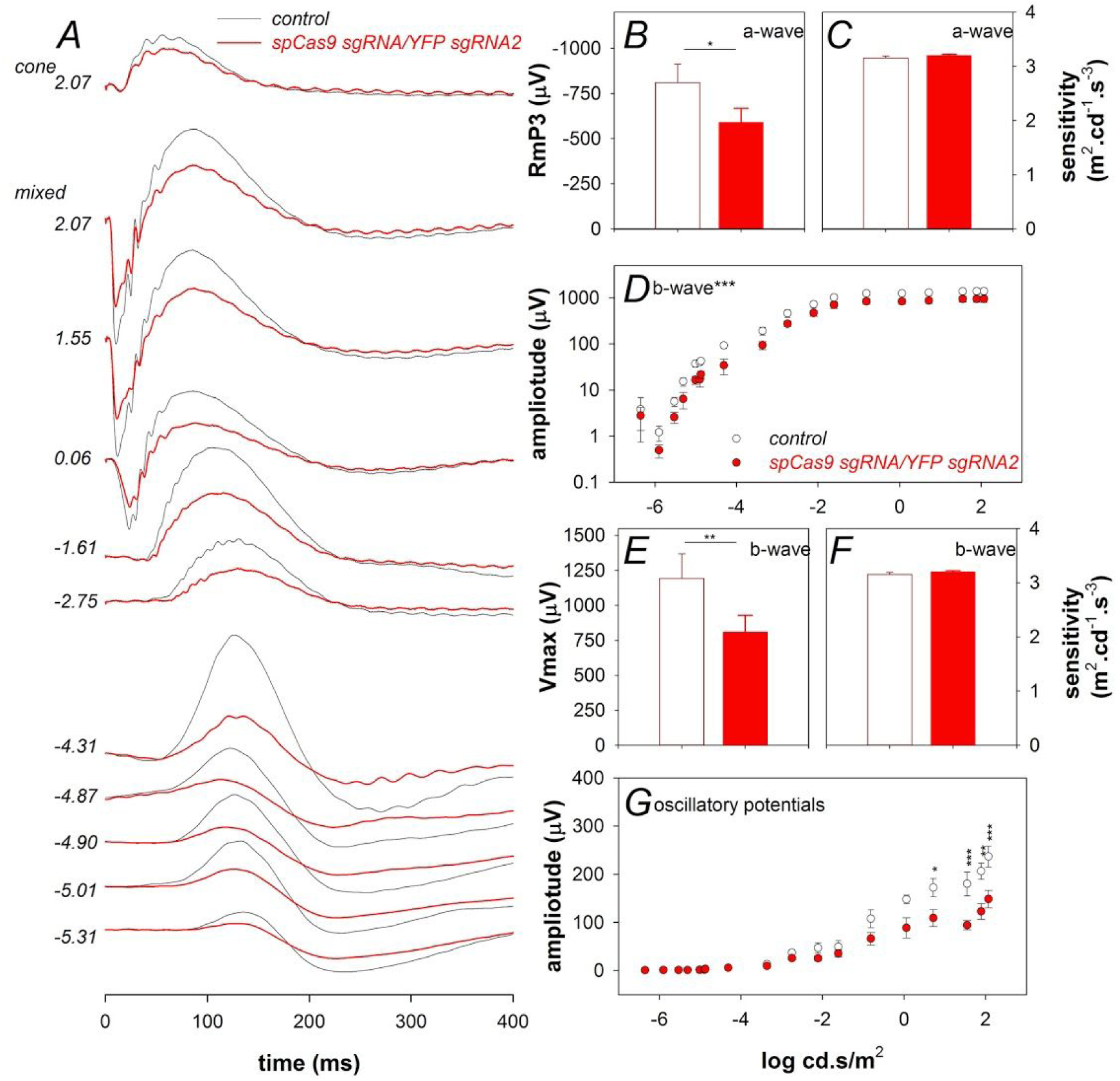
SpCas9 sgRNA/YFP sgRNA decreased retinal function. (A) Averaged ERG waveforms at selected intensities for control (n=4, black) and treated eyes (n=4, red). (B) Groups average photoreceptoral (a-wave) saturated amplitude for contralateral control (unfilled) and treated eyes (filled). (C) Photoreceptoral sensitivity to light. (D) Intensity response characteristics across the entire range of intensities. (E) Bipolar cell amplitude. (F) Bipolar cell sensitivity to light. (G) Inner retinal amacrine cell mediated response (oscillatory potentials). Data are expressed as the Mean ± SEM. Statistical analysis between groups was performed using two-tailed Student’s t-test. Asterisks denotes significance *P<0.05, **P<0.001, ***P<0.0001.

**Supplementary Figure 4.**
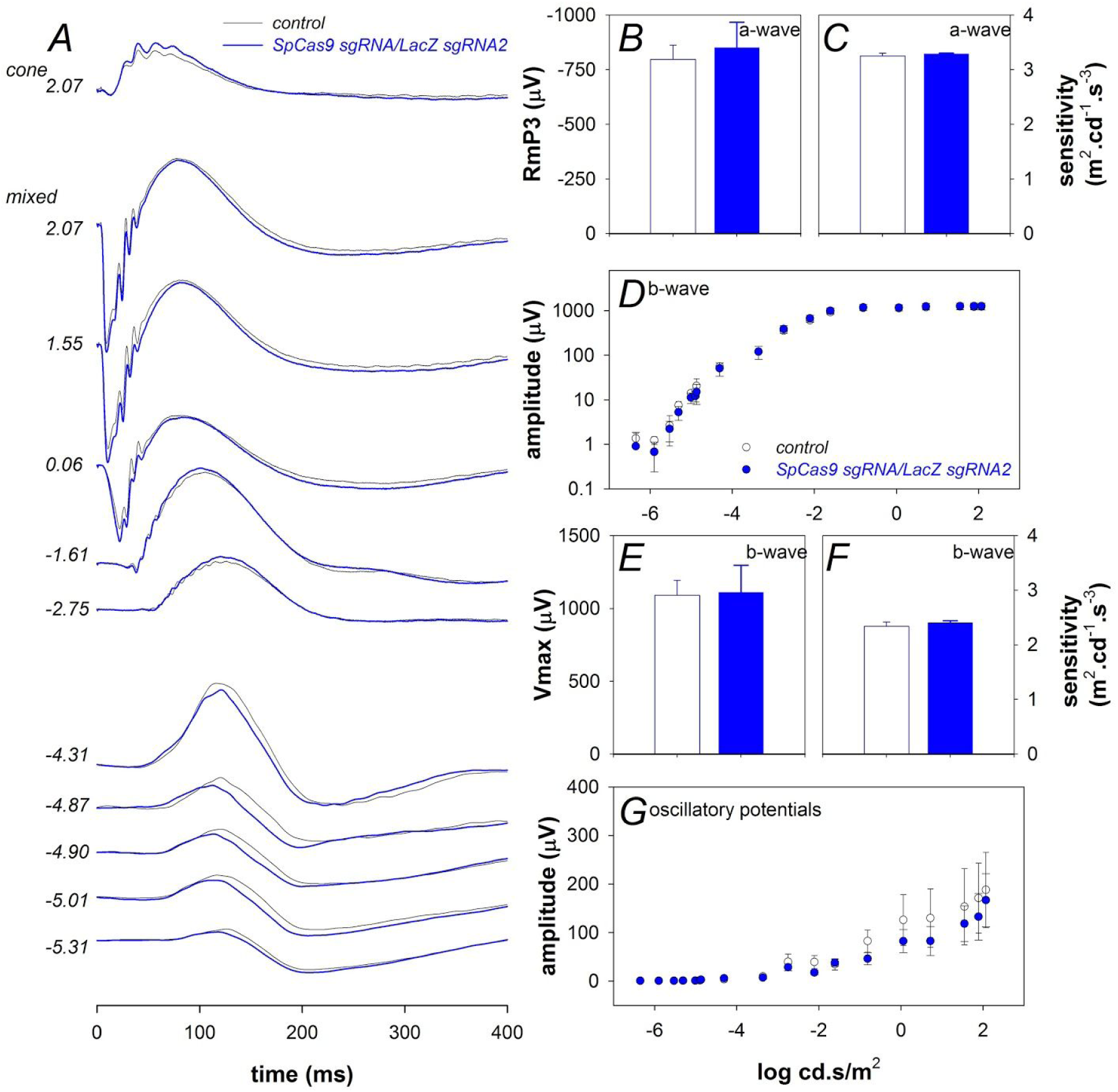
SpCas9 sgRNA/LacZ sgRNA does not affect retinal function. (A) Averaged ERG waveforms at selected intensities for control (n=3, black) and treated eyes (n=3, blue). (B) Groups average (±SEM) photoreceptoral (a-wave) saturated amplitude for contralateral control (unfilled) and treated eyes (filled). (C) Photoreceptoral sensitivity to light. (D) Intensity response characteristics across the entire range of intensities. (E) Bipolar cell amplitude. (F) Bipolar cell sensitivity to light. (G) Inner retinal amacrine cell mediated response (oscillatory potentials). Data are expressed as the Mean ± SEM. Statistical analysis between groups was performed using two-tailed Student’s t-test.

**Supplementary Figure 5.**
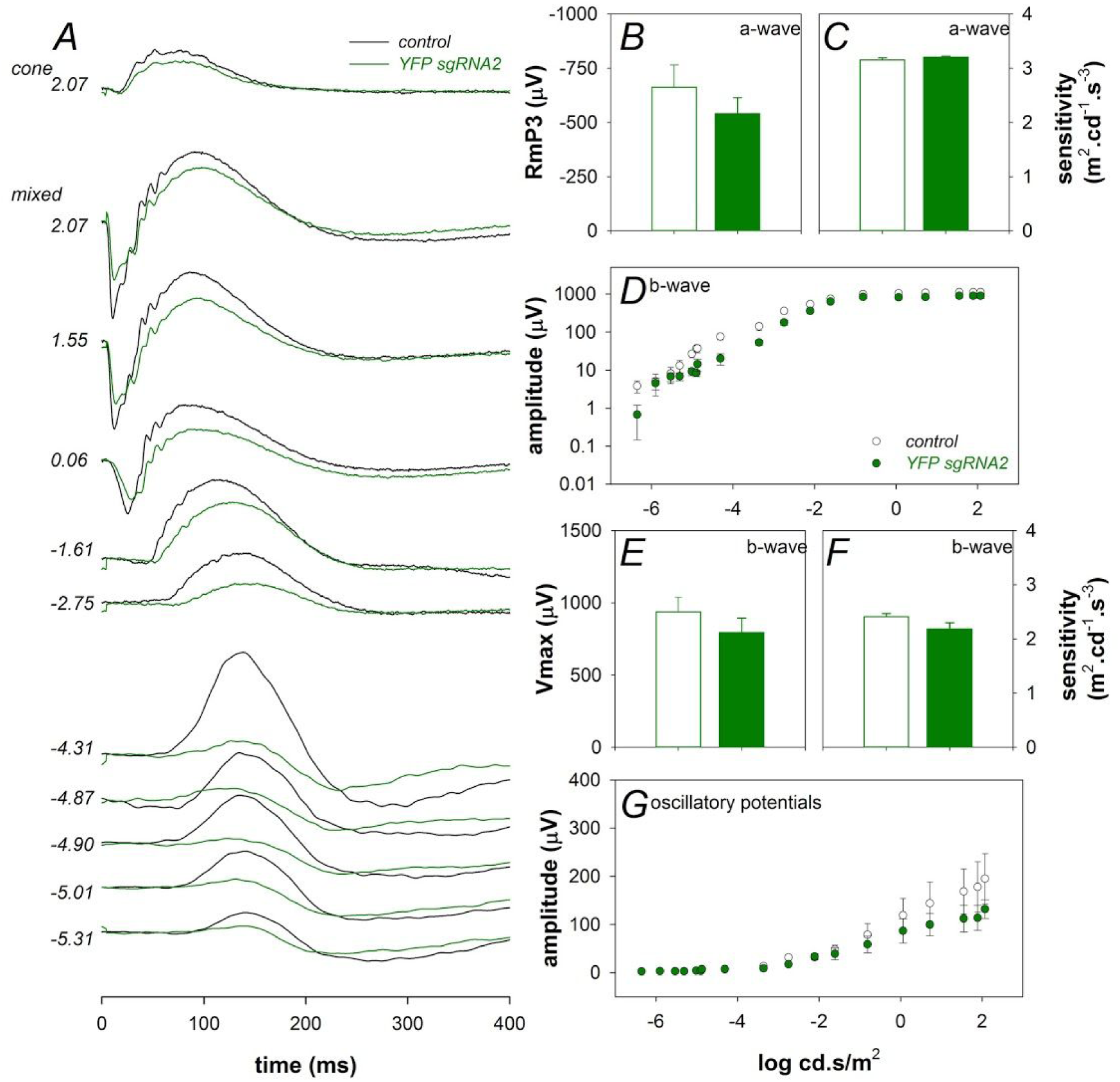
YFP sgRNA2 alone affects inner retinal function. (A) Averaged ERG waveforms at selected intensities for control (n=5, black) and treated eyes (n=5, green). (B) Groups average (±SEM) photoreceptoral (a-wave) saturated amplitude for contralateral control (unfilled) and treated eyes (filled). (C) Photoreceptoral sensitivity to light. (D) Intensity response characteristics across the entire range of intensities. (E) Bipolar cell amplitude. (F) Bipolar cell sensitivity to light. (G) Inner retinal amacrine cell mediated response (oscillatory potentials). Data are expressed as the Mean ± SEM. Statistical analysis between groups was performed using using two-tailed Student’s t-test.

**Supplementary Figure 6.**
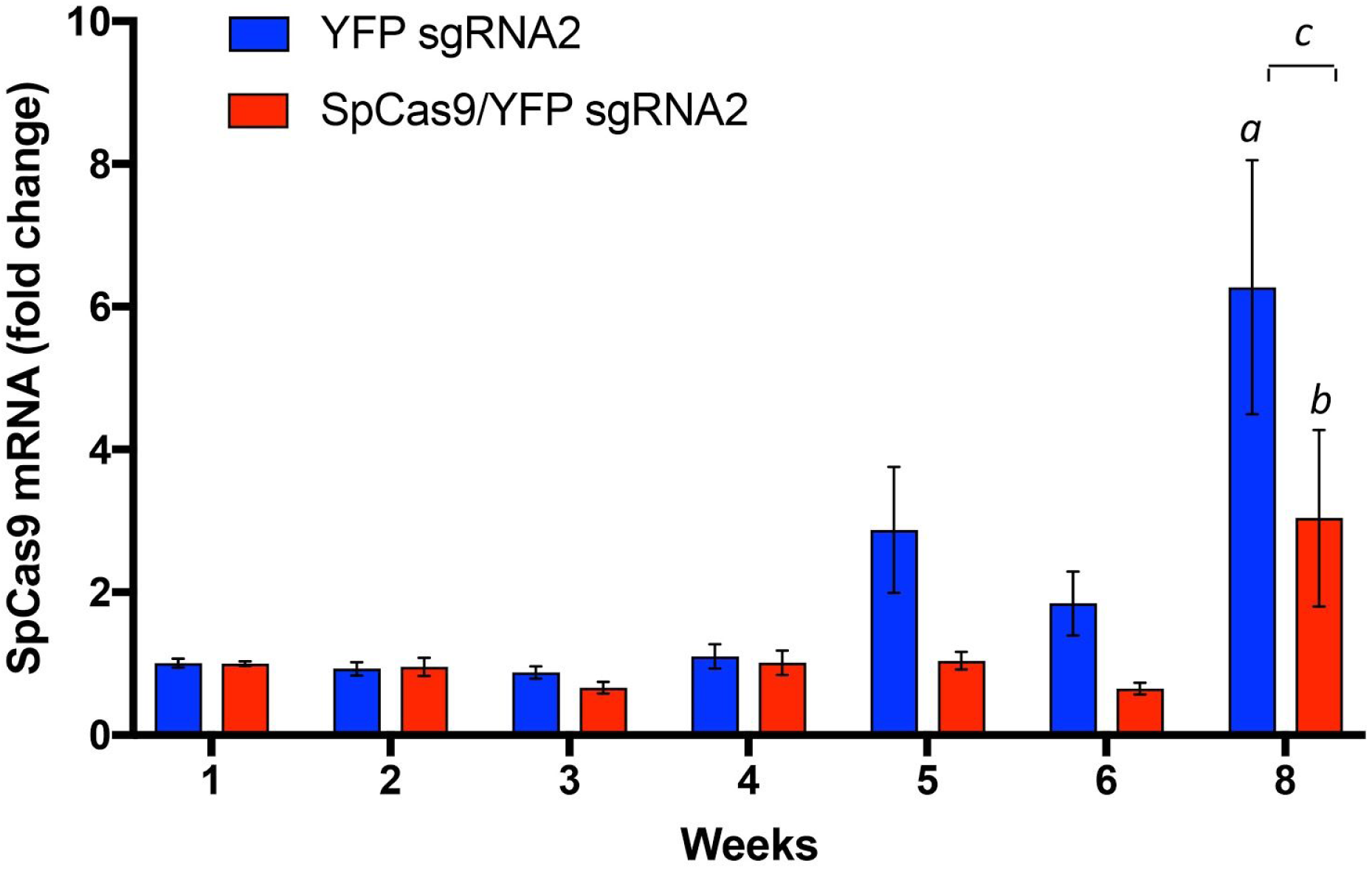
Time course of SpCas9 mRNA expression in the mouse retina. SpCas9 mRNA were isolated from the mice retina administrated with AAV2-SpCas9 sgRNA/YFP sgRNA2 or AAV2-YFP sgRNA2 at 5, 6 and 8 weeks after intravitreal injection. Relative fold change of SpCas9 expression was normalized by week 1 from each treatment group. Representative data are shown for 5-6 retinas per group/time point and expressed as Mean ± SEM. Statistical analysis between groups was performed using Two-way ANOVA followed by Sidak's multiple comparisons test, *a*, YFP sgRNA2: 1 vs 8 weeks, p=0.002. *b*, SpCas9/YFP sgRNA: 1 vs 8 weeks, *p*=0.7043. *c*, YFP sgRNA2 vs SpCas9/YFP sgRNA, p=0.0142.

**Supplementary Figure 7.**
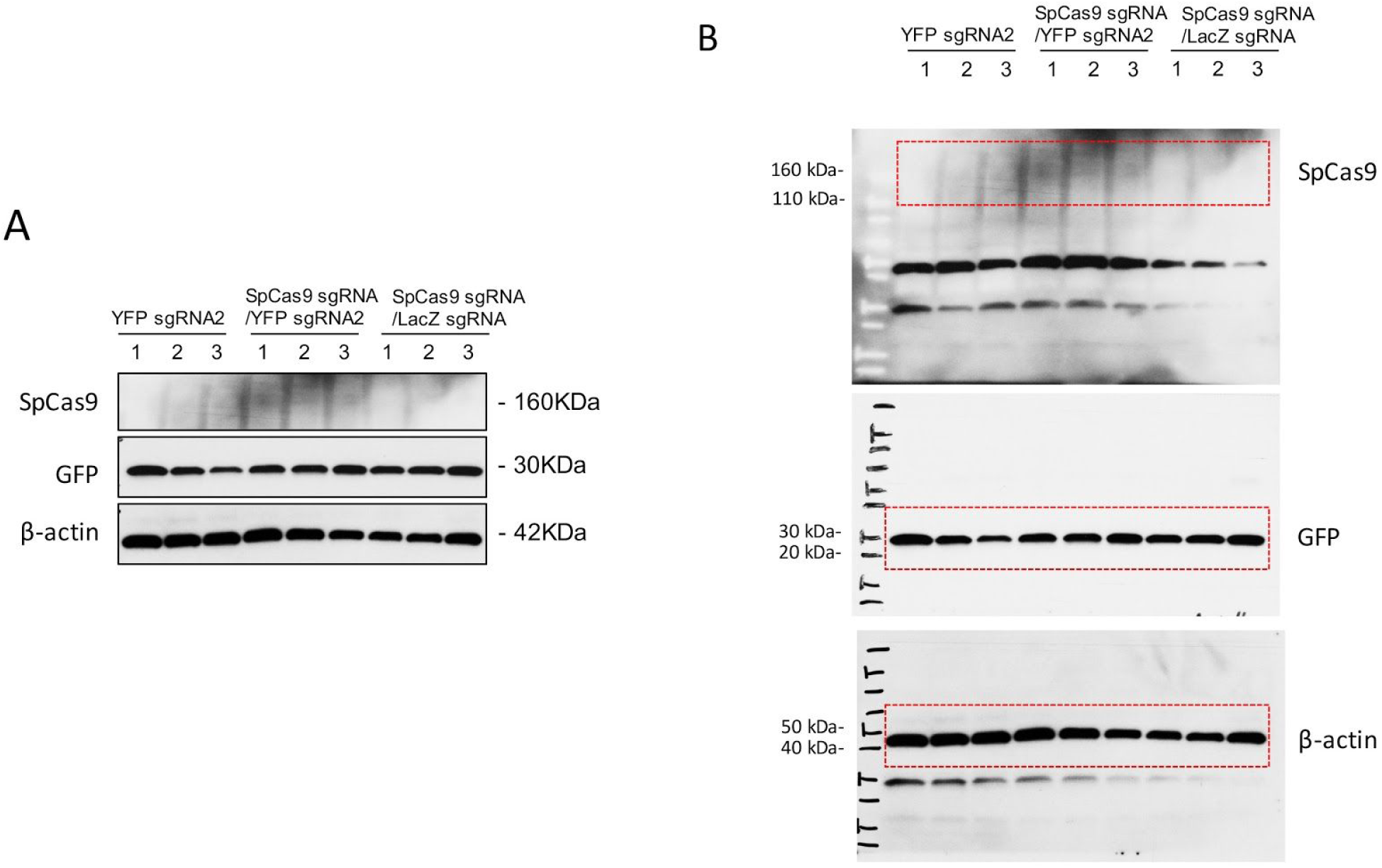
Representative western blots of retinal SpCas9 expression. (A) SpCas9 and YFP expression in the mouse retinas were assessed by western blot analysis. (B) Uncropped western blot images.Three retinas from different mice in each group were randomly selected for western blot analysis.

**Supplementary Figure 8.**
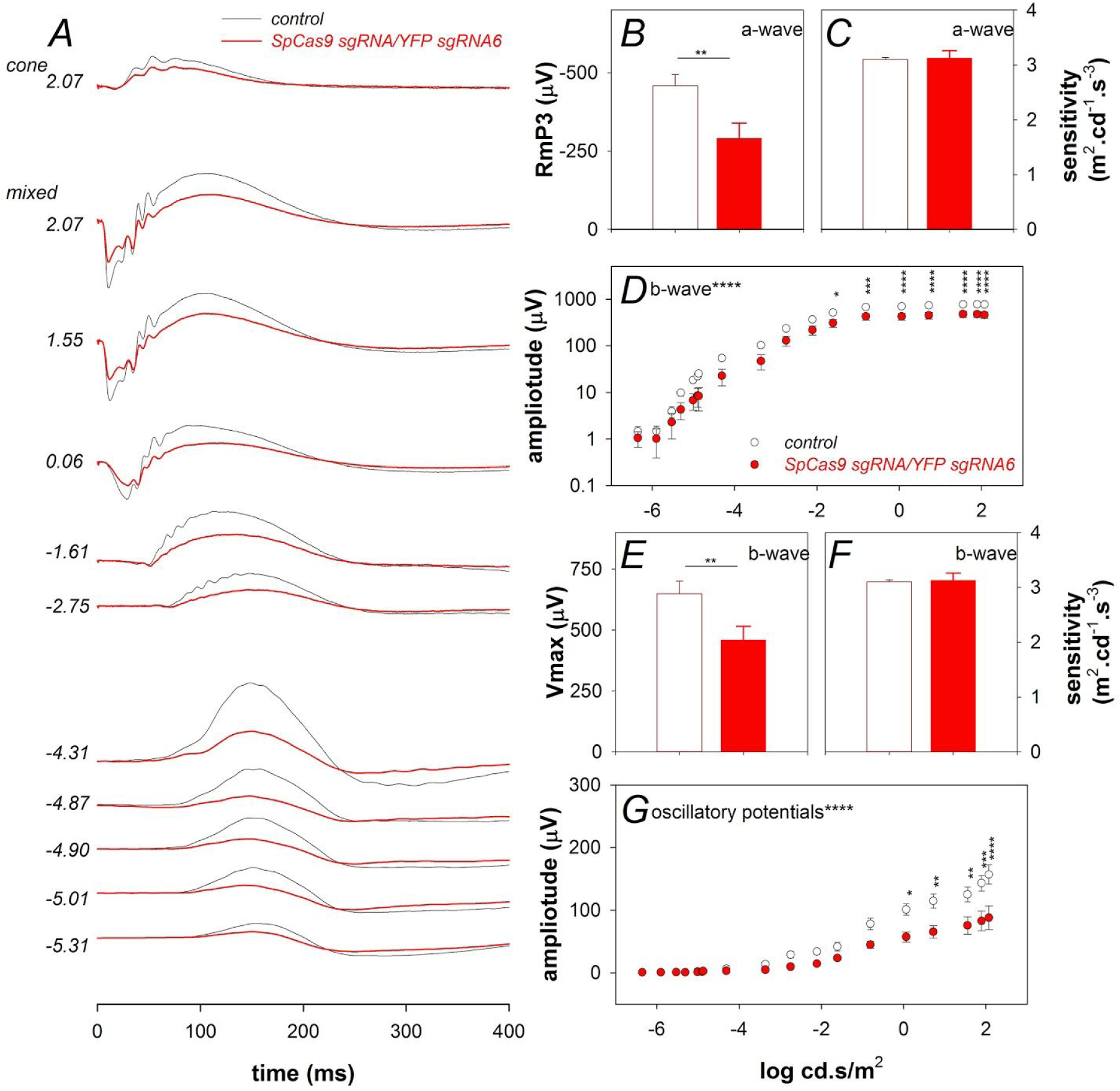
SpCas9 sgRNA/YFP sgRNA6 decreased retinal function. (A) Averaged ERG waveforms at selected intensities for control (n=10, black) and treated eyes (n=10, red). (B) Groups average (±SEM) photoreceptoral (a-wave) saturated amplitude for contralateral control (unfilled) and treated eyes (filled). (C) Photoreceptoral sensitivity to light. (D) Intensity response characteristics across the entire range of intensities. (E) Bipolar cell amplitude. (F) Bipolar cell sensitivity to light. (G) Inner retinal amacrine cell mediated response (oscillatory potentials). Data are expressed as the Mean ± SEM. Statistical analysis between groups was performed using two-tailed Student’s t-test. Asterisks denotes significance *P<0.05, **P<0.001, ***P<0.0001, ****P<0.00001.

**Supplementary Figure 9.**
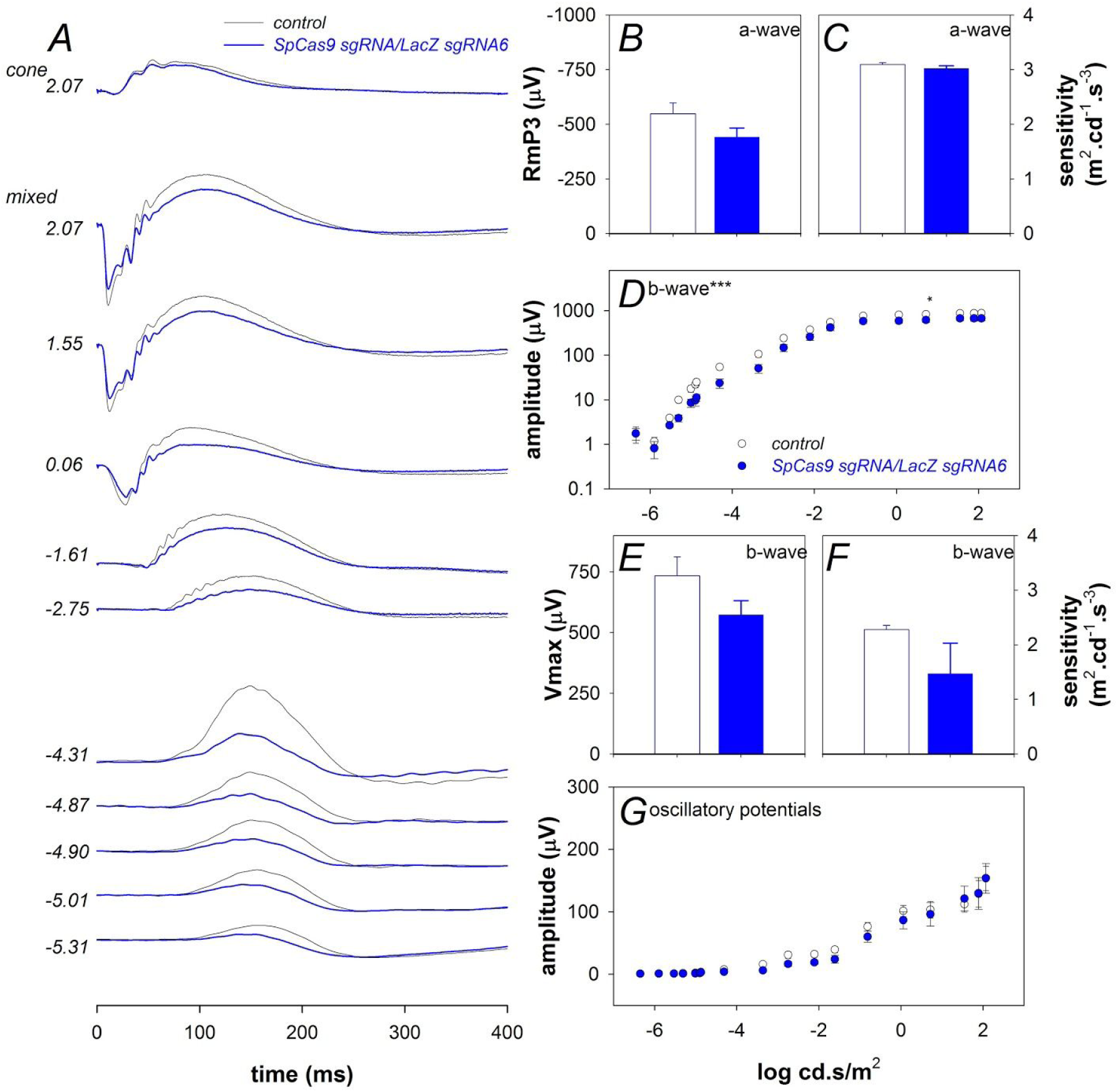
SpCas9 sgRNA/LacZ sgRNA6 does not affect retinal function. (A) Averaged ERG waveforms at selected intensities for control (n=8, black) and treated eyes (n=8, blue). (B) Groups average photoreceptoral (a-wave) saturated amplitude for contralateral control (unfilled) and treated eyes (filled). (C) Photoreceptoral sensitivity to light. (D) Intensity response characteristics across the entire range of intensities. (E) Bipolar cell amplitude. (F) Bipolar cell sensitivity to light. (G) Inner retinal amacrine cell mediated response (oscillatory potentials). Data are expressed as the Mean ± SEM. Statistical analysis between groups was performed using two-tailed Student’s t-test. *P<0.05.

**Supplementary Figure 10.**
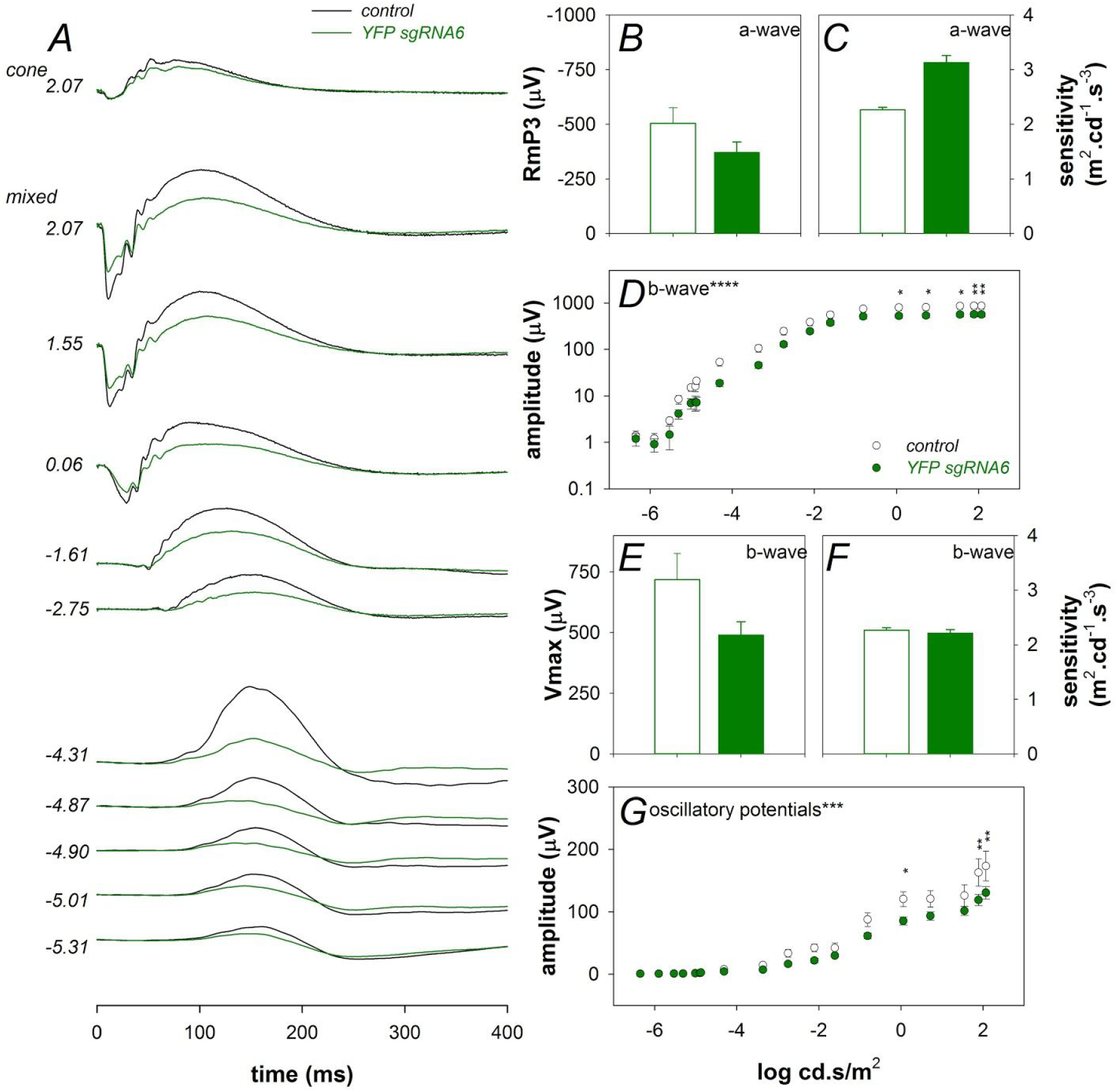
YFP sgRNA6 alone affects inner retinal function. (A) Averaged ERG waveforms at selected intensities for control (n=8, black) and treated eyes (n=8, green). (B) Groups average photoreceptoral (a-wave) saturated amplitude for contralateral control (unfilled) and treated eyes (filled). (C) Photoreceptoral sensitivity to light. (D) Intensity response characteristics across the entire range of intensities. (E) Bipolar cell amplitude. (F) Bipolar cell sensitivity to light. (G) Inner retinal amacrine cell mediated response (oscillatory potentials). Data are expressed as the Mean ± SEM. Statistical analysis between groups was performed using two-tailed Student’s t-test. Asterisks denotes significance. *P<0.05, **P<0.001.

